# Grassland irrigation and fertilisation alter vegetation height and vegetation within temperature and negatively affect orthopteran populations

**DOI:** 10.1101/2020.01.17.910042

**Authors:** Jean-Yves Humbert, Sarah Delley, Raphaël Arlettaz

**Affiliations:** Division of Conservation Biology, Institute of Ecology and Evolution, University of Bern, Baltzerstrasse 6, 3012 Bern, Switzerland

**Keywords:** agriculture, Alps, arthropods, conservation, grasshoppers, meadow

## Abstract

European mountain meadows are hosting an exceptionally rich biodiversity. While they have long been exposed to land abandonment, they are nowadays additionally threatened by agriculture intensification through aerial irrigation and slurry application. The consequences of this intensification on arthropods are not well documented and studies are needed to fulfil this knowledge gap. Six experimental management treatments combining a full factorial design and a gradual level of fertilisation and irrigation were implemented in 2010 in twelve different montane and subalpine Swiss meadows. In 2013, orthopterans were sampled to assess the influence of the management practices on their population. In addition changes in vegetation height and temperature induced by intensification were recorded in order to better appraise underlying mechanisms. Intensification had a negative impact on Caelifera (grasshoppers); with decreases of up to 70% in densities and 50% in species richness in the most intensively managed treatment plots. In parallel intensification induced an increase in mean vegetation height and a cooling of up to 4.2 °C (10 cm aboveground) within most intensively managed plots. These microhabitat and microclimate changes are likely to have affected Caelifera development, in particular thermophilous species. In contrast, Ensifera (bush crickets) densities and species richness did not respond to the management treatments. The use of irrigation (without fertilisation) had limited impacts on orthopterans and microclimate. In conclusion, orthopterans, in particular Caelifera, are relatively sensitive to grassland intensification and to conserve the full community, mountain agricultural systems need to maintain extensively managed meadows.

## 1. Introduction

In Europe, mountain meadows represent one of last remnant of exceptionally diverse semi-natural grassland types (Veen et al., 2009). However two new management practices are spreading in alpine regions and threaten these biodiversity rich habitats: irrigation with sprinkler and fertilisation with liquid manure (Maurer et al., 2006; Riedener et al., 2013). Both of these practices modify the vegetation community and structure which in turn affects the arthropod populations (Andrey et al., 2014; Schwab et al., 2002). Arthropods play an important role in grassland systems and beyond; they provide or at least participate in a range of ecosystem services such as pollination, decomposition process or pest control (e.g. Sutter and Albrecht, 2016) and are primordial food items for many vertebrates (e.g. Arlettaz, 1996; Wilson et al., 1999). This underlines the importance to preserve their abundance and diversity. So far fertilisation has been shown to have a negative impact on arthropod species richness ensuing from a reduction of vegetation diversity (Haddad et al., 2009; Haddad et al., 2000). On the other hand it seems to boost herbivore abundances through an increase in plant tissue nitrogen and to have cascading effect on other arthropods functional groups (Andrey et al., 2016; Haddad et al., 2001; Hudewenz et al., 2012). In contrast the effects of irrigation on arthropods remain poorly documented. Therefore current knowledge does not allow determining the irrigation and fertilisation thresholds that should not be exceeded in order to maintain a functional and diverse arthropod community in mountain meadows (but see Andrey et al., 2016; Lessard-Therrien et al., 2018).

The goal of the present study was to assess the response of orthopteran species richness and density to gradual levels of fertilisation and irrigation in montane and subalpine meadows. In these grasslands, orthopterans represent the most important insect group in term of biomass (Blumer and Diemer, 1996). They are a key component of the diet of many insectivorous species and an important decline in their density would have cascading effect on higher trophic level (Britschgi et al., 2006; Vickery et al., 2001). In addition, orthopterans are recognized bioindicators for grasslands as they readily respond to management changes (Buri et al., 2013) and are sensitive to a set of vegetation parameters (Le Provost et al., 2017). First, orthopterans are sensitive to microclimate (Löffler and Fartmann, 2017) which varies with vegetation height and density (Song et al., 2013). As ectotherms organisms, their development rate, body size, reproductive success and other physiological processes are depending on temperature. Each species has its own thermal sensitivity: for example eurythermal species such as *Pseudochorthippus parallelus* are very tolerant and can adapt to a range of microclimatic conditions while thermophilous species such as *Stenobothrus lineatus* are restricted to warm and dry habitats (van Wingerden et al., 1991; Willott and Hassall, 1998). Microclimatic conditions influence thus the orthopteran community. Second, the habitat diversity hypothesis stipulates that more diverse a habitat is the more species it is likely to host (Báldi, 2008). At an orthopteran scale microhabitat diversity is function of the vegetation structural heterogeneity which is to some extent correlated with plant diversity (e.g. Morris, 2000; Woodcock et al., 2009) and vegetation height (Andrey et al., 2014). Finally food availability is a limiting factor for the expansion of any organism. A sufficient proportion of grass is essential to maintain Caelifera (grasshoppers) density as they almost exclusively feed on it (Ibanez et al., 2013). Ensifera (bush crickets) on their side have a more diversified diet composed of small invertebrates and grasses and are thus less dependent on specific food resources (Baur et al., 2006). The first aim of the present study was to investigate how orthopteran populations respond to gradual level of irrigation and fertilisation and to determine whether an optimum management intensity maximising both density and species richness exists. The second aim was to measure the changes in vegetation height and aboveground temperatures induced by intensification, and to determine whether orthopteran responses can be explained by these changes.

In the short term fertilisation has been shown to increase vegetation structure and phytomass production (Andrey et al., 2014) while in the long term it induces a loss of plant species richness and a homogenisation of the vegetation cover (e.g. Lessard-Therrien et al., 2017; Marini et al., 2008; Socher et al., 2013). In addition it usually induces a shift in plant community toward higher percentage of grass and legumes (Rudmann-Maurer et al., 2008; Socher et al., 2013) whereas irrigation favours grass species and increase nitrogen (N) mineralization by plants (Jeangros and Bertola, 2000; Riedener et al., 2013). Finally both inputs boost the productivity and thus create a denser and taller sward (Bassin et al., 2012; Marini et al., 2009). Consequently we expected aboveground temperature to gradually cool down along the intensification gradient (Song et al., 2013). We expected orthopteran densities to increase at mid-intensity as a consequence of better food quality (Hudewenz et al., 2012; Joern et al., 2012) and to decrease in highly intensified plots due to the detrimental effect of microclimate cooling (van Wingerden et al., 1992). Concerning orthopterans species richness, we expected it to decrease steadily along the intensification gradient due to the disappearance of thermophilous species and the loss of microhabitats (Fournier et al., 2017; van Wingerden et al., 1991).

## 2. Material and methods

### 2.1 Study sites

The study was carried out in the canton of Valais, an inner Alps valley of Switzerland which experiences a continental climate with cold and wet winter and dry and hot summers: mean annual temperature amounts 10.7°C and mean annual precipitation achieves 517 mm (2000 ‒ 2014 mean in Sion, 482 m a.s.l.). In 2010, twelve extensively managed meadows were selected within this region; they were situated between 790 and 1740 m a.s.l (Appendix 1).

### 2.2 Experimental design

In 2010, within each meadows (n = 12), six different management treatments were randomly allocated to plots of 20 m in diameter spaced from each other by at least 5 m. The first plot served as a control (C). The second and third plots were only irrigated (I) or fertilised (F), and the fourth to sixth plots were irrigated and fertilised (I+F; Table 1). The exact amount of fertiliser applied at each site depended on the theoretical maximum hay yield achievable locally with two harvests per year, calculated using pre-experimental hay yield and site elevation (for details see Appendix A in Andrey et al., 2016). Accordingly, sites were divided in three groups; 1, 2 and 3, where I+F 3/3-plots received, respectively, 40, 60 or 80 [kg N·ha-1·yr-1]. Within group, mid-intensive (F and I+F 2/3) and low-intensive (I+F 1/3) plots, received respectively two-thirds and one-third of the maximum fertilisation dose. It has been decided to follow these prescriptions in order to obtain results within realistic agronomical systems. Fertiliser consisted of organic dried manure NPK pellets (MEOC SA, 1906 Charrat, Switzerland), and mineral potassium oxide (K_2_O) dissolved in water to reach the equivalent of standard-farm liquid manure consisting namely of 2.4 kg of usable nitrogen, 2 kg of phosphate (P_2_O_5_), and 8 kg of potassium oxide (K_2_O) per m^3^ of solution. Every year the plots were fertilised once in early spring and once after the first cut (June or July), each time half of the annual fertiliser amount was applied (except for the 1/3-plots that were fertilised only once in spring). Treatments I and I + F were additionally irrigated weekly from mid-May to end of August, except when heavy rainfall occurred (≥20 mm over the previous week). Irrigation thresholds were chosen on the basis of Calame Calame et al. (1992) experiment. Accordingly I and I+F 2/3 matched the recommendation for the best profitability of water input (20 mm/week) while low-intensive (I+F 1/3) and high-intensive (I+F 3/3) management treatments received respectively half and one and an half of this dose (Table 1).

**Table 1:** Management practices applied to the six different experimental treatment plots. Abbreviations for treatments: C = control; F = fertilised; I = irrigated; F+I 1/3, F+I 2/3 and F+I 3/3 = fertilised and irrigated at respectively 1/3, 2/3 or 3/3 of the maximum dose. The exact 3/3 dose of fertiliser applied at each site followed the management norm recommended to achieve maximum hay yield at a given locality, and were classed in three groups. Note that I and F, received the same amount of water or fertiliser as I+F 2/3.

| Management treatment | Mowing regime [no. of cut-yr<sup>-1</sup>] | Irrigation [mm-week<sup>-1</sup>] | Fertilisation [kg N·ha<sup>-1</sup>·yr<sup>-1</sup>] |  |  |
| --- | --- | --- | --- | --- | --- |
|  |  |  | Group 1 | Group 2 | Group 3 |
| C | 1 | 0 | 0.0 | 0.0 | 0.0 |
| I | 2 | 20 | 0.0 | 0.0 | 0.0 |
| F | 2 | 0 | 53.3 | 40.0 | 26.6 |
| I+F 1/3 | 2 | 10 | 26.6 | 20.0 | 13.3 |
| I+F 2/3 | 2 | 20 | 53.3 | 40.0 | 26.6 |
| I+F 3/3 | 2 | 30 | 80.0 | 60.0 | 40.0 |

### 2.3 Orthopterans sampling

Orthopterans were sampled in 2013 with a biocenometer (open trap) made of a net fastened around a strong circular wire so as to provide a total capture area of 1 m^2^ (as described in Humbert et al., 2012). Two sampling sessions were performed: one shortly before the first cut (between 12 June and 12 July) and one 4–6 weeks after it (between 13 August and 31 August). The date at which meadows were sampled was function of their elevation. During both sessions, eight samples were regularly taken per treatment plot. All the individuals trapped within the biocenometer were caught and identified on site. Adults were identified to species level while juveniles were classified into suborders (Caelifera or Ensifera). All samplings were done on sunny days between 10 am and 5 pm.

### 2.4 Vegetation height record

Vegetation height was measured as the average vegetation stratum height in a 10 cm radius around a meterstick. Eight records were taken per plot during the orthopterans samplings and averaged to have one value per plot. All the measurements were performed by the same person.

### 2.5 Temperature record

To record aboveground temperatures I-buttons DS1921G-F Thermochron (Maxim Integrated Products/Dallas) were used, which are self-sufficient systems measuring and recording temperature in 0.5 °C increment. In each plot one of those device was randomly placed at 5 m from the centre, fixed on a stick 10 cm above the ground. I-buttons recorded temperature hourly from beginning of May to end of August. They were removed shortly before the first cut and replaced within a few days. The data from the ten days following mowing event were removed from the analysis to reduce noise due to the manipulation of the devices. Average daily temperature was calculated as the mean temperature between 12 am and 4 pm while average nocturnal temperature was calculated as the mean temperature between 12 pm and 4 am.

### 2.6 Statistical analyses

Treatments effects were analysed with linear mixed models (LMMs) or generalized linear mixed models (GLMMs) using the *lmer*, respectively *glmer*, functions from the *lme4* package for R (Bates et al., 2015). Response variables were orthopteran densities, species richness, vegetation height and temperatures; they were analysed with either Poisson (Caelifera and Ensifera densities) or Gaussian (others) distribution. Though, vegetation height had to be log-transformed in order to achieve normal distribution of residuals. The fixed effects were the treatments (C, I, F, I+F 1/3, I+F 2/3, I+F 3/3), and the random intercept effects were the study sites in all the analyses. When using the Gaussian distribution, p-values were obtained using the *lmerTest* package Kuznetsova, 2017 #775}. Caelifera and Ensifera responses were analysed separately as they differ in their ecology (Baur et al., 2006). Vegetation height, temperature and density data were analysed per sampling session while species data were pooled. Models always fulfilled model assumptions, notably residuals normal distribution and homoscedasticity.

Structural equation modelling (SEM), using the *lavaan* package (Rosseel, 2012), were further used to determine if fertilisation and irrigation influence orthopterans directly or/and indirectly through changes in vegetation height or aboveground temperature. In this SEM analysis, the water and slurry inputs were treated as two continuous variables with four levels: control with no input = 0; I+F 1/3 = 1; I, F and I+F 2/3 = 2; and I+F 3/3 = 3. As a first step, a set of candidate models was developed. Candidate models always included slurry and water inputs as two independent variables, and then their effects on orthopterans were either direct, indirect through vegetation height or aboveground temperature, or both. In addition, the number of paths was set to maximum four which led to a total of twenty candidate models (see Appendix 2 for a graphical representation of all SEM candidate models). Second, all models were ran and kept only if the overall fit of the respective model was satisfactory. To assess model fit, the chi-square test (if *P* > 0.05), the comparative fit index CFI (if CFI > 0.95), the root mean square error of approximation RMSEA (if RMSEA < 0.07) and the standardised root mean square residuals SRMR (if SRMR < 0.08) were used (Hooper et al. 2008). Third, retained models were ranked based on AIC values (Akaike information criterion) and the model with the lowest AIC plus the model(s) within a Δ AIC < 2 were considered the most plausible model(s). If several models were ranked within a Δ AIC < 2, the model with the highest R-square was chosen as best model. The *lavaan.survey* package, which applies robust maximum likelihood method to estimate the standard errors, was used to include the meadows as random effect in the SEM (Oberski, 2014). All statistics were performed using R version 3.6.1 (R Core Team, 2019).

## 3. Results

Due to unfortunate field circumstances, orthopteran densities could not be sampled in site Cordona before mowing. Similarly, no aboveground temperatures were recorded in Eison and Grimentz after mowing. Therefore all related analyses were based on n = 11 or n = 10 sites respectively.

### 3.1 Orthopterans density

Mean density of orthopterans varied greatly among meadows and plots. It ranged from 0.13 to 24.38 individuals per m^2^ during the first sampling session and from 0.65 to 27.38 individuals per m^2^ during the second sampling session. Treatments were found to have significant effects on Caelifera densities while none were detected on Ensifera. Note that low densities of Ensifera limited the power of the analysis on this suborder.

Before mowing the highest Caelifera densities were found within C-plots, mean ± standard error (SE) = 8.42 ± 2.73, that hosted ∼30–40% more individuals than I-plots (5.68 ± 2.05, *P* = 0.016) and F-plots (4.77 ± 1.82, *P* < 0.001) and >70% more individuals than I+F 1/3-plots (2.43 ± 0.66, *P* < 0.001), I+F 2/3-plots (2.02 ± 0.87, *P* < 0.001) and I+F 3/3-plots (2.26 ± 0.81, *P* < 0.001; see Fig. 1a and Appendix 3 for detailed model outputs). Concerning the Ensifera, the highest densities were found within F-plots (0.94 ± 0.16) and the lowest within I+F 3/3-plots (0.45 ± 0.14), though differences were not statistically significant (Fig. 1b and Appendix 3). C-plots (0.66 ± 0.18), I-plots (0.80 ± 0.16), I+F 1/3-plots (0.53 ± 0.23) and I+F 2/3-plots (0.57 ± 0.17) had intermediate densities. After mowing Caelifera and Ensifera densities did not differed across treatments (Fig. 1c, Fig. 1d and Appendix 3).

**Fig. 1.**
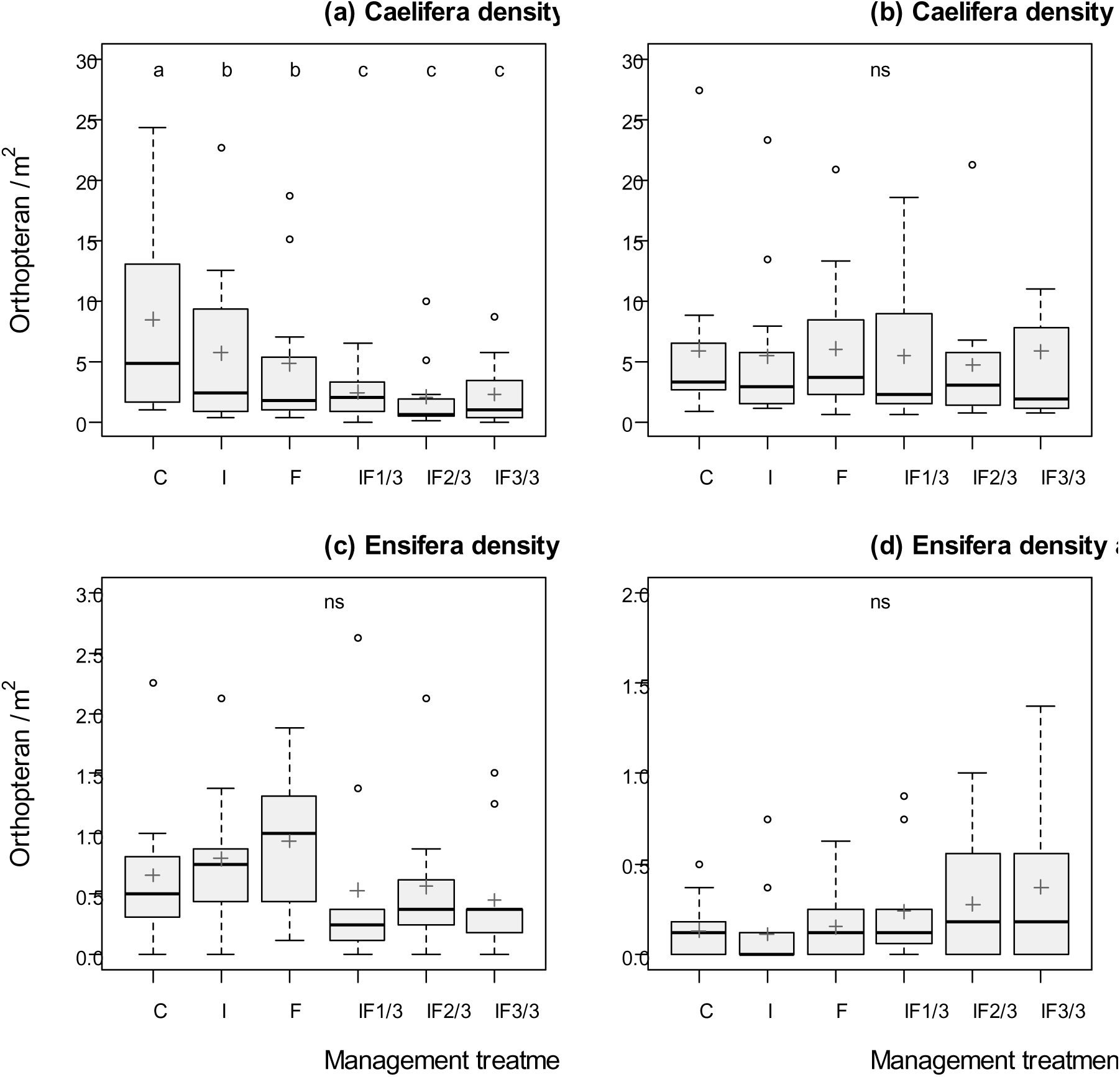
Responses of orthopteran densities (individuals / m^2^) to the six different management treatments: a) Caelifera densities before mowing; b) Caelifera densities after mowing (note that in the IF3/3-plot a point at 31.9 individuals / m^2^ do not appear on the figure); c) Ensifera densities before mowing; d) Ensifera densities after mowing. Abbreviations for treatments: C = control; F = fertilisation only; I = irrigation only, IF1/3 = irrigation and fertilisation at low dose, IF2/3 = irrigation and fertilisation at medium dose, IF3/3 = irrigation and fertilisation at high dose. Different letters indicate significant differences between treatments at an alpha rejection value set to 0.05. Bold lines represent medians, cross the means; boxes the first and third quantiles.

### 3.2 Orthopteran species richness

A total of 21 species was recorded within all plots, seven of which were Ensifera and fourteen of which were Caelifera (see Appendix 1 for detailed list). The minimum number of species found within a plot was one and the maximum was nine. Management practices significantly affected the Caelifera species richness. The highest Caelifera species richness was found within C-plots (4.6 ± 0.5), that hosted similar species number than I-plots (4.25 ± 0.5) and F-plots (4.2 ± 0.5) but ∼30% more species than I+F 1/3-plots (3.2 ± 0.5, *P* < 0.001) and I+F 2/3-plots (3.5 ± 0.5, *P* = 0.004) and 50% more species than I+F 3/3-plots (2.3 ± 0.4, *P* < 0.001; see Fig. 2a and Appendix 4 for detailed model outputs). Contrariwise, no significant effects were detected on the Ensifera species richness (see Fig. 2b and Appendix 4).

**Fig. 2.**
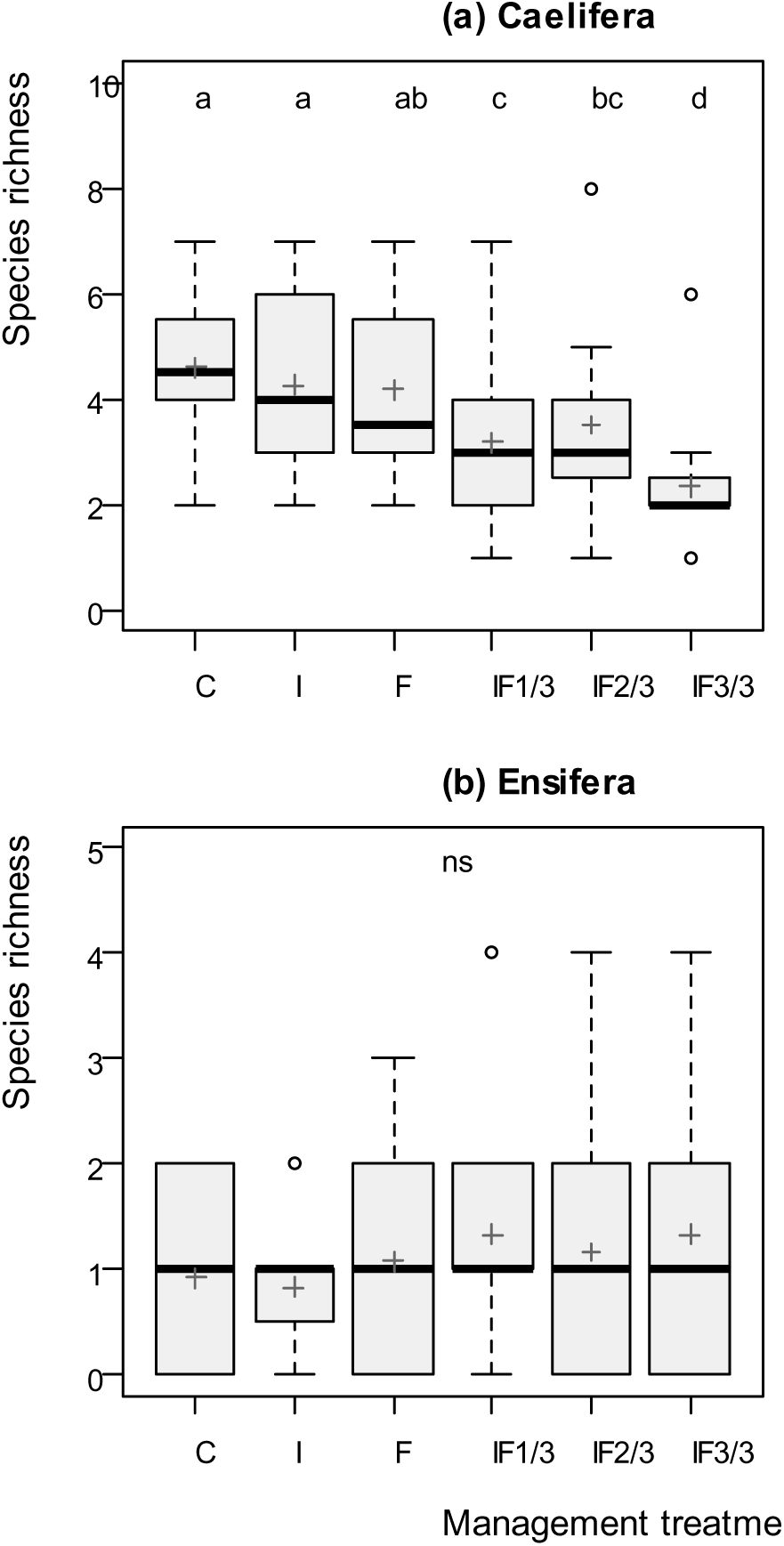
Responses of Caelifera (a) and Ensifera (b) species richness to the six different management treatments. For treatment abbreviations and boxplot descriptions see legend of figure 1.

### 3.3 Vegetation height

Before mowing vegetation stratum height was the tallest in I+F 3/3-plots (61.8 cm ± 3.5 cm), it was slightly shorter in I+F 1/3-plots (52.1 ± 7.7, *P* = 0.016), I+F 2/3-plots (51.9 ± 2.3, *P* = 0.080) and F-plots (49.9 ± 3.4, *P* = 0.023) while it grew half less in C-plots (31.8 ± 2.9, *P* < 0.001) and I-plots (36.9 ± 3.1, *P* < 0.001; see Fig. 3a and Appendix 5 for detailed model outputs). After mowing the same trend was observed with tallest sward found within I+F 3/3-plots (24.2 ± 2.0), followed by I+F 2/3-plots (18.3 ± 2.2, *P* = 0.006) and then I+F 1/3-plots (14.0 ± 2.2, *P* < 0.001), F-plots (12.8 ± 2.2, *P* < 0.001) and I-plots (12.3 ± 1.2, *P* < 0.001), while C-plots vegetation (9.0 ± 2.2, *P* < 0.001) was more than twice shorter (Fig. 3b and Appendix 5).

**Fig. 3.**
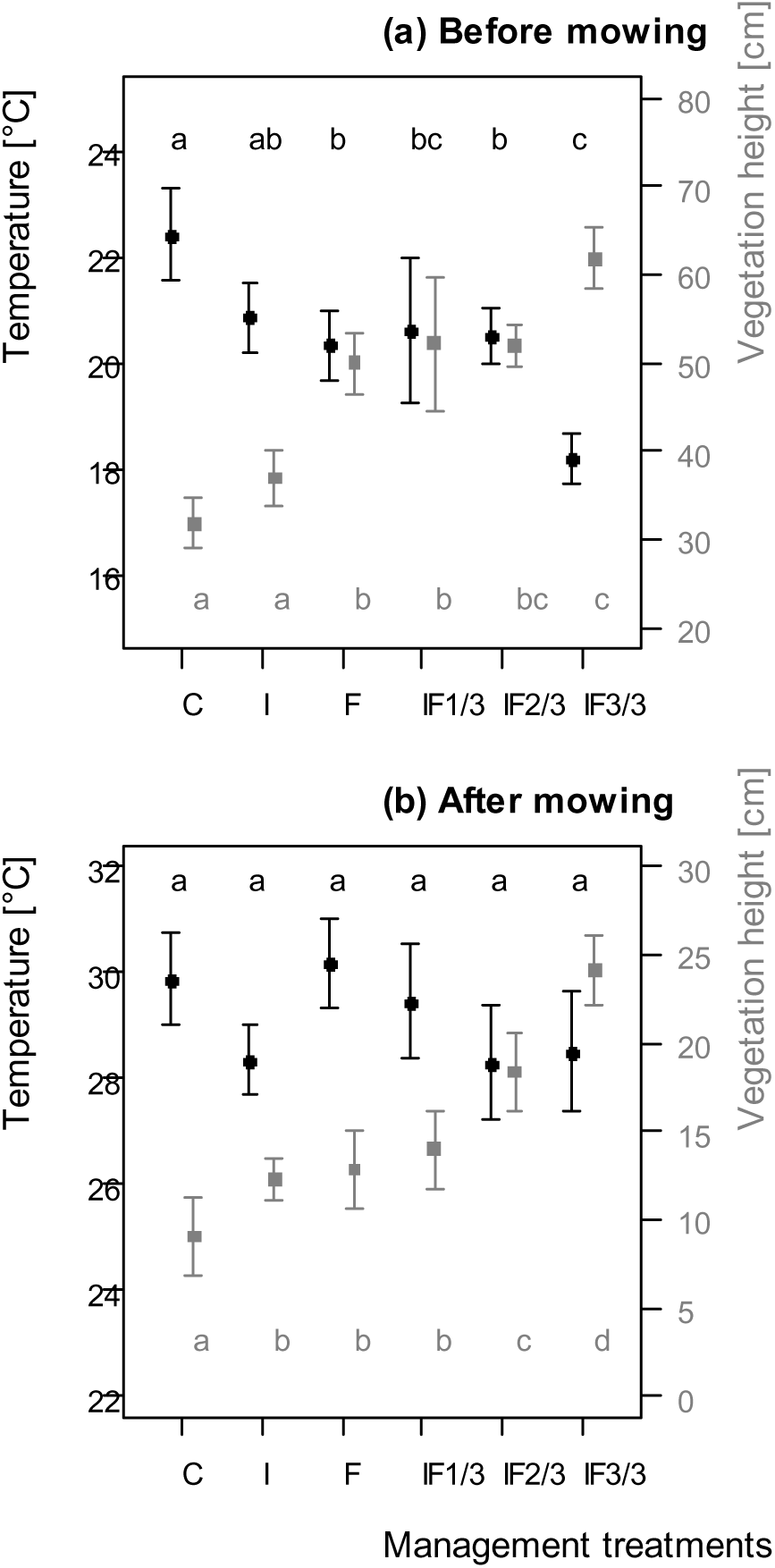
Aboveground diurnal temperature (black circles) and vegetation height (grey squares) according to the six different management treatments, before (a) and after mowing (b). Mean values ± SE of the raw data are shown. Different letters indicate significant differences between treatments at an alpha rejection value set to 0.05. For treatment abbreviations see legend of figure 1.

### 3.4 Temperature

Before mowing, mean diurnal aboveground temperature was the warmest in C-plots (22.4 ± 0.9 °C), then temperatures in I-plots (20.9 ± 0.7), I+F 1/3-plots (20.6 ± 1.4), I+F 2/3-plots (20.5 ± 0.6) and F-plots (20.3 ± 0.7) were 1.5–2.1 °C colder (all *P* < 0.05 except for I-plots) while it was over 4.2 °C colder in I+F 3/3-plots than in C-plot (18.2 ± 0.5, *P* < 0.001). See Fig. 3a and Appendix 6 for detailed model outputs. Given the noticeable negative relationship between temperatures and vegetation heights, an additional LMM was run between both (Fig. 4; Estimate = −3.383, SE = 0.894, *P* < 0.001).

**Fig. 4.**
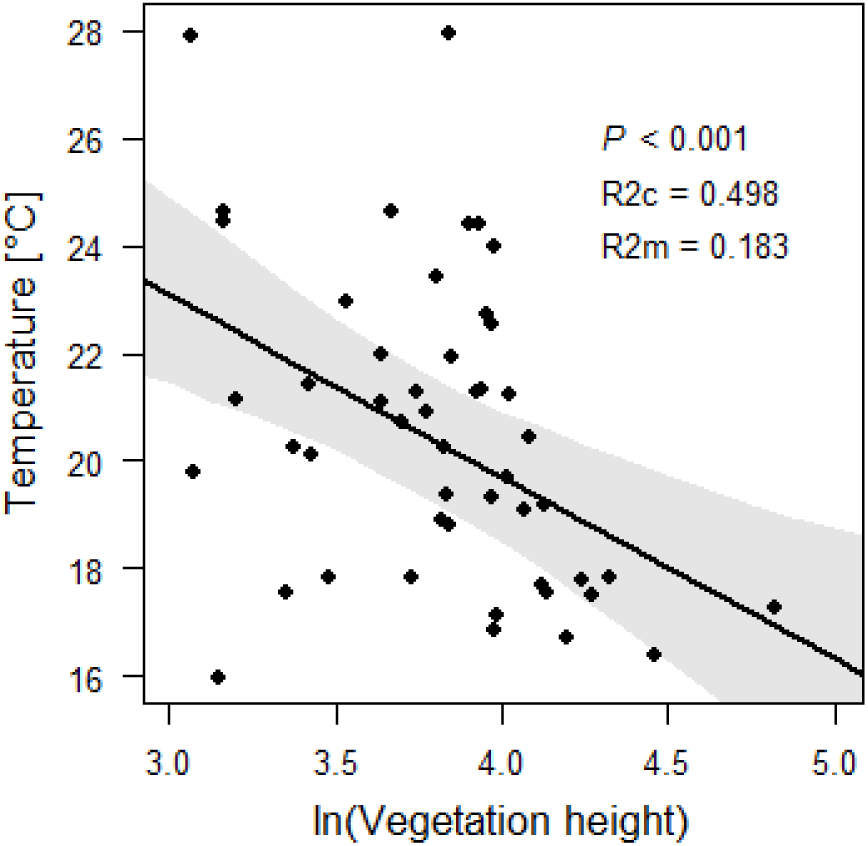
Negative relationship between aboveground diurnal temperature and log-transformed vegetation height (in cm) before mowing. The regression line is drawn from the LMM outputs with 95% confidence intervals. Marginal r^2^ (R2m) represents the percentage of variance explained by the fixed effects only, whereas conditional r^2^ (R2c) is the percentage explained by both fixed and random effects grouped (Nakagawa and Schielzeth, 2013).

After mowing, diurnal aboveground temperature was the highest in C-plots (29.9 ± 0.9) and F-plots (30.2 ± 0.9). I+F 1/3-plots (29.5 ± 1.2), I+F 3/3-plots (28.5 ± 1.4), I-plots (28.4 ± 0.7) and I+F 2/3-plots (28.3 ± 1.1) were respectively 0.4 to 1.7 °C colder than C-plots, but differences were not statistically significant (Fig. 3b and Appendix 6). Treatments did affect nocturnal temperatures but differences were not biologically relevant (in order of 0.1–0.2 °C) and are thus not further discussed.

### 3.5 Structural equation modeling (SEM)

As treatments did not affect Ensifera densities nor species richness SEMs were run only on Caelifera. Before mowing the best SEM model (chi-square = 0.084, d.f. = 1, *P* = 0.772; CFI = 1; RMSEA < 0.001; SRMR = 0.006) explaining changes in Caelifera densities included both indirect effects of slurry and water inputs through vegetation height plus a direct effect of water input (Fig. 5a). The best SEM models explaining Caelifera densities after mowing (chi-square = 0.612, d.f. = 1, *P* = 0.434; CFI = 1; RMSEA < 0.001; SRMR = 0.013) and Caelifera species richness (chi-square = 0.612, d.f. = 1, *P* = 0.434; CFI = 1; RMSEA < 0.001; SRMR = 0.014) were the same. They included both direct effects of slurry and water inputs plus an indirect effect of water input through aboveground temperature measured after mowing (Fig. 5b and 5c). However, for the density after mowing, only one path was statistically significant; i.e. the effect of water input on aboveground temperature. For species richness the direct effect of slurry input on Caelifera species richness was significant, as well as the effect of water input on aboveground temperature coupled with a non-significant effect (P = 0.253) of aboveground temperature on Caelifera species richness. For Caelifera species richness, all combination of SEM models with vegetation height and aboveground temperature measured before and after the first mowing were tried.

**Fig. 5.**
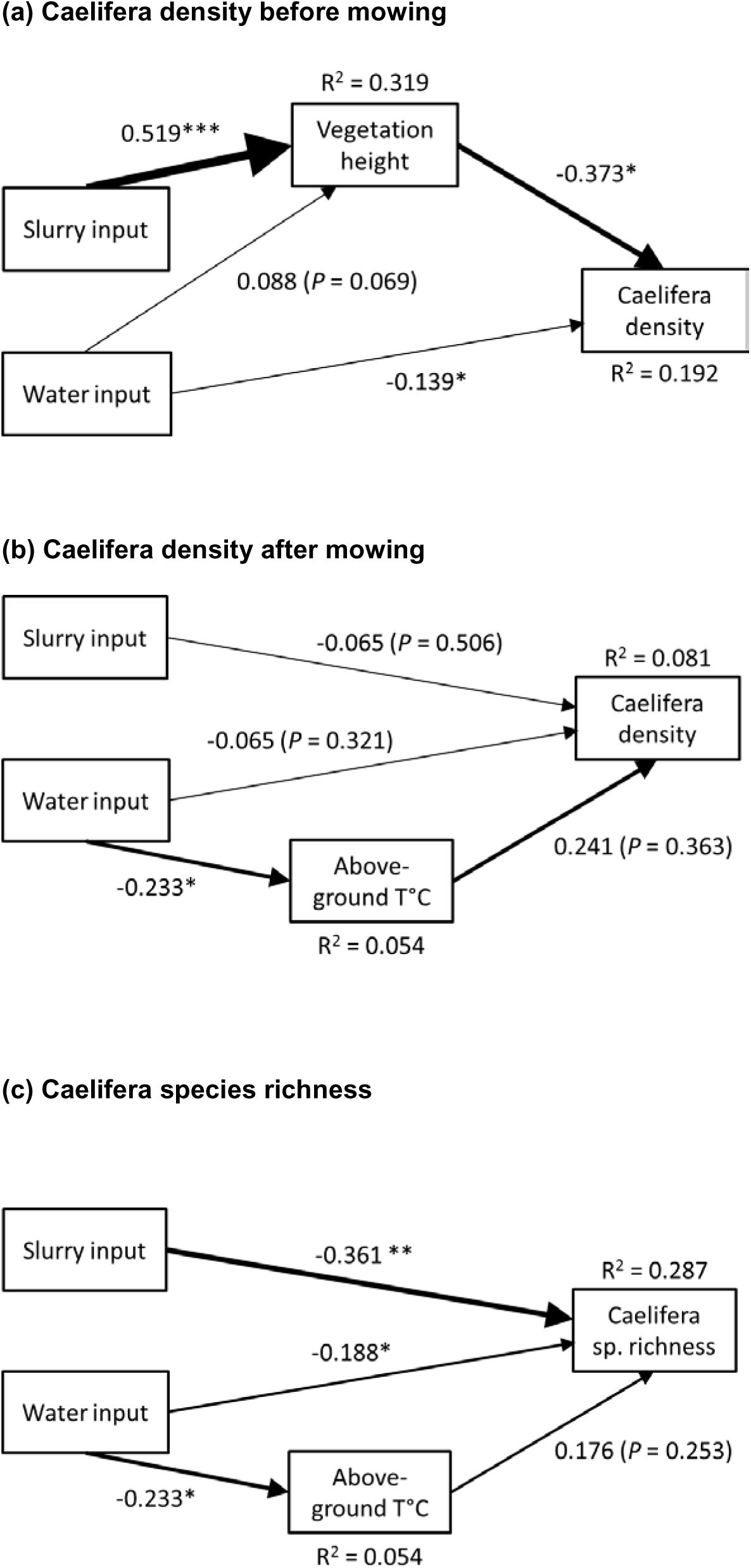
Best structural equation model (SEM) explaining the influences of slurry and water inputs on: (a) Caelifera density before mowing; (b) Caelifera density after mowing; and (c) Caelifera species richness. Standardized path coefficients are shown beside each path, with the level of statistical significance indicated by asterisks (\**P* < 0.05; \*\**P* < 0.01; \*\*\**P* < 0.001). The width of the arrows depicts the strength of the effect and R^2^ values represent the proportion of variance explained for each dependent variable. All candidate SEM models can be found in Appendix 2.

## 4. Discussion

This study shows that mountain grassland fertilisation and irrigation combined, greatly affects Caelifera (grasshoppers) by decreasing their densities and species richness, but that the use of irrigation only (at two-thirds of the maximum dose) has relatively limited impacts. On the other hand, overall Ensifera (bush crickets) densities and species richness did not respond to the management treatments. This study also demonstrates that intensification practices induce an important drop in aboveground temperatures parallel to an increase in vegetation heights. Microclimate cooling has often been suggested as a potential mechanism to explain responses of various arthropod groups to intensification (e.g. Gardiner et al., 2002; Löffler and Fartmann, 2017; Marini et al., 2009), but the link had never been clearly demonstrated.

In the following subsections we first present the effects of the management practices on vegetation and microclimate. We then state its effects on orthopteran density and species richness and discuss how these are linked to vegetation height and microclimate. Finally we discuss the conservation implications of the present study.

### 4.1 Effects on vegetation and microclimate

Combined irrigation and fertilisation led to twice taller swards in the most intensively managed plots (I+F 3/3) compared to control plots, which was expected as water and nitrogen are limiting factor for vegetation growth in dry mountain region (Bassin et al., 2012; Tasser and Tappeiner, 2002). Before the first cut, irrigation alone had lesser effect, likely as a consequence of the wet spring of 2013. However after the cut both inputs had an equivalent positive effect on plants regrowth and their combination amplified their respective effects.

Aboveground temperatures were linked to vegetation height: the taller the sward became the less sunlight reached the ground and the less it was warmed (see also Song et al., 2013). Consequently the temperatures differences between the most intensively managed plots and the controls were of up to 4.2°C at 10 cm aboveground. After the cut, vegetation regrew progressively which reduced the surface temperature differences among plots. Vegetation height is not the only factor explaining microclimate; other parameters such as vegetation density and canopy cover do influence quantity of sunshine reaching the soil and thus indirectly temperature (van Wingerden et al., 1992), as well as sprinkler irrigation as demonstrated here. This explains why surface temperatures varied more along the intensification gradient than swards heights.

### 4.2 Effects on orthopteran densities

Different responses were observed between the two suborders. Before mowing Caelifera densities were respectively 30 to 40% lower in plots that had been either irrigated or fertilised. The combination of both inputs was even worse leading to over 70% reduction in density regardless of quantities applied (i.e. densities were divided by three). Our SEM indicates that this negative effect is due to a combination of a direct negative effect of water input and an indirect effect of slurry input through vegetation height. As most individuals were low-mobile larvae during the first sampling session their distribution reflects their birth place. Therefore reasons for differences could be one or a combination of following factors: 1) females favoured lower sward sites to oviposit; 2) larval development was altered and survival rate was lower in plots with tall vegetation (Willott and Hassall, 1998); or 3) eggs hatching was delayed in taller vegetation and had not occurred yet at the time plots were sampled, which is detrimental to population as it reduces individuals chance to complete their life cycle and to reproduce (van Wingerden et al., 1991; Weiss et al., 2013). Management intensification did not impact Ensifera densities. This might reflect the fact that in average Ensifera emerged earlier in the season compared to Caelifera, when vegetation height and thus microclimate differences were less pronounced among management practices. Moreover their development is globally less dependent on temperatures (Bieringer and Zulka, 2003).

The mowing event may have dispersed the individuals all over the meadow area (Humbert et al., 2012) so the densities of Caelifera found during the second sampling session were not directly related to number found during the first sampling session. Nevertheless, we observed that generalist species such as *P. parallelus* dispersed more or less evenly across the plots while the specialized thermophilous species such as *S. lineatus* or *Omocestus haemorrhoidales* recolonized the warmest plots. A boom in Caelifera eggs hatching probably occurred in the days following mowing due to warmer microclimate as about half of the individuals sampled during the second session were still at larval stage. After mowing Ensifera were slightly, though not statistically, more numerous in more intensively managed plots and there were virtually no more larvae. This pattern was due to the preponderant presence of adults *Tettigonia viridissima* and *Roeseliana roeselii* that favoured the tall swards found in intensified plots. However densities remained very low with less than 0.5 Ensifera per m^2^.

We had hypothesized that at moderate management intensity level orthopteran densities would be maximized, benefiting from increased food supply without being impacted by microclimate changes (Hudewenz et al., 2012). However, results are not in accordance with our hypothesis. Overall results seem to indicate that in the investigated mountain meadows food resource is not a limiting factor for Caelifera while temperature might be.

### 4.3 Effects on orthopteran species richness

Highest Caelifera species richness was found in the control plots and was maintained in plots that were either irrigated or fertilised but the combination of both inputs had detrimental effects: species loss reached 50% in the most intensive plots, i.e. richness was divided by two. This trend is in line with previous observational studies done in the Alps or Prealps (Marini et al., 2008; Schlegel and Schnetzler, 2018), as well as conclusions form studies carried on in lowland regions (e.g. Chisté et al., 2016; Knop et al., 2006) and it confirms the general detrimental effects of grassland management intensification on orthopteran species richness.

The SEM indicates strong direct negative effects of slurry and water input, plus an indirect effect of water input through changes in aboveground temperature, though the last path between aboveground temperature and Caelifera species richness was not statistically significant. The direct effect of slurry and water input should not be interpreted as strictly direct, it could also be indirect through variables not included in the model. It is known that in addition to aboveground temperature, orthopteran community composition is related to several other parameters such as vegetation structural heterogeneity (Jerrentrup et al., 2014), percentage of bare ground (Weiss et al., 2013), plant species composition (Fournier et al., 2017; Gardiner et al., 2002; Ibanez et al., 2013), and management regime (e.g. our control plots were cut once a year while irrigated and fertilised plots twice; see also Buri et al., 2013). The parallel drop in species richness and temperatures suggests that either thermophile species chose deliberately not to oviposit within more intensive plots – showing a cumulative effect from previous year – or the eggs laid within colder plots poorly developed and larvae never reached maturity (Willott and Hassall, 1998). Willott and Hassall (1998) showed that a difference of 5°C in ambient temperature – air temperature differences reached 4.2°C in our case – considerably affects Caelifera fitness: most sensitive species experiments a development time 50% longer, a reduction of 25% in body mass and a drop of 50% in pods production. Altogether, our findings support the hypothesis that the microclimate conditions within intensively managed plots became too cold for thermophilous species. Though, we do not claim that it is the only explanatory mechanism. *De facto* temperature changes do not explain the relatively lower Caelifera species richness found within low-intensity plots (I+F 1/3) compared to irrigated or fertilised only plots.

Contrariwise to Caelifera, Ensifera species richness was not affected by intensification. This suborder is known to be less sensitive to microclimate than Caelifera and to depend more on vegetation structure (Baur et al., 2006; Bieringer and Zulka, 2003). However, a change in community composition accompanying intensification was noticed. Large species such as *T. viridissima* or generalists such as *R. roeselii* favouring tall vegetation which offers good singing spots and shelter (Baur et al., 2006; Buri et al., 2013) were more often found in intensively managed plots compared to extensively managed plots. In the contrary *Plactycleis albopunctata* or *Decticus verrucivorus* which are species associated with warm and dry habitat (Baur et al., 2006) were occasionally found in control plots, but not within intensively managed ones. This reflects the ecological diversity of Ensifera and might explain why their global species richness remained stable among experimental plots. It has to be noted that the power of the analyses was constrained due to relatively low Ensifera species richness (the average ± standard deviation was 1.1 ± 1.0 across all plots).

### 4.4 Conclusions and conservation implications

Aerial irrigation and fertilisation with liquid manure are two novel management practices currently spreading in dry alpine regions (Riedener et al., 2013). While on one hand these practices benefit biodiversity in the sense that by increasing grass yield (Andrey et al., 2014; Bassin et al., 2012) they support continuity of local farming so as to keep montane and subalpine semi-natural grasslands open (Riedener et al., 2014; Rudmann-Maurer et al., 2008), on the other hand they become a threat to biodiversity when too much inputs are applied (this study).

In contrast to observational studies, the experimental approach adopted in this study has the advantage that measured differences between treatments were not affected (biased) by environmental parameters such as soil, elevation or surrounding landscape, neither by climate or past management history. In addition, results are based on controlled, quantitatively based, levels of grassland irrigation and fertilisation that are related to what is done in the practice. On the other hand it has some limitations due to the size and proximity of the plots that could have blurred the signal. Despite the close proximity of the experimental plots, very clear evidences that intensification lower both Caelifera densities and species richness were found, which are then conservative findings.

Results demonstrate that the use of irrigation input alone has moderate impacts on orthopterans and on microclimate while the combination of irrigation and fertilisation inputs is harmful to orthopterans even at low dose. The first finding is good news for all stakeholders (including farmers, conservationists and policy-makers) as irrigation alone can increase hay yield without affecting biodiversity (see also Jeangros and Bertola, 2000; Riedener et al., 2013) and is allowed in subsidized extensively managed meadows registered under Swiss agri-environment schemes. On the other hand fertilisation alone and the combination of both inputs must be forbidden in grasslands where conservation of invertebrate and vertebrate fauna is of concern as a drop in grassland invertebrate densities can have dramatic bottom-up effects on higher trophic levels (Britschgi et al., 2006). Management intensification had even stronger impacts on Caelifera densities than on species richness as detrimental effects were already visible in low intensity plots. Combine with our precedent findings on plants (REF (Andrey et al., 2014; Lessard-Therrien et al., 2017), bryophytes (Boch et al., 2018), Auchenorrhyncha (Andrey et al., 2016), spiders and ground beetles (Lessard-Therrien et al., 2018), this study darken the optimist perspective to find an intermediate management intensity threshold that provides decent agronomical yield and that has only limited negative effects on biodiversity.

The second important contribution of this study is the quantitative assessment of changes in microhabitat and -climate induced by intensification. The effective cooling (more than 4 °C at 10 cm above ground) measured exceeded expectations and can have huge impacts on the development of local micro-fauna (Logan et al., 2006). While we acknowledge that we only prove a causal connection between changes in vegetation height and Caelifera densities before mowing, aboveground temperature was retained (regardless of its non-statistically significant last path) in the best SEMs to explain Caelifera density after mowing and species richness, and the models showed very good fitting properties. In addition there was a strong negative correlation between vegetation height and sward temperature which highlights that both effects can hardly be separated.

## 5. Acknowledgment

We would like to thank A. Hayoz-Andrey for help implementing the study, S. Mettaz, M. Clerc, G. Delley, S. Laribi, P. Buri, J. Flury, A. Spang, D. Schlunke and J. Coflica for field assistance, our colleagues at the Division of Conservation Biology, University of Bern, as well as the members of the project accompanying group. We are also thankful to the farmers for their collaboration. This project was financially supported by the Cantons of Graubünden and Valais, by the Swiss National Science Foundation (grants 31003A_125398 and 31003A 149656 to RA) and the Swiss Federal Offices for Agriculture and the Environment.

## Appendices

## Appendix 1.

Location name of all study sites (n = 12), geographic coordinates (WGS84), elevation, sampling date, and number of orthopterans of each species caught during the two sampling sessions (i.e. before and after mowing). In addition, meadows were classified in three groups according to the maximum hay productivity potential of the site (see Material and Methods section for more details). Data are missing for the first session in Cordona due to technical problems.

See Supplemental File.

## Appendix 2.

Structural equation modelling (SEM) were used to determine if fertilisation and irrigation influence orthopterans directly or/and indirectly through changes in vegetation height or aboveground temperature. The chart represents the full structural equation model including all potential paths. However, the number of paths of the candidate models was set to maximum four and they always included slurry and water inputs which led to a total of twenty candidate models. SEMs were ran on Caelifera densities before and after mowing and on Caelifera species richness (pooled sampling session).

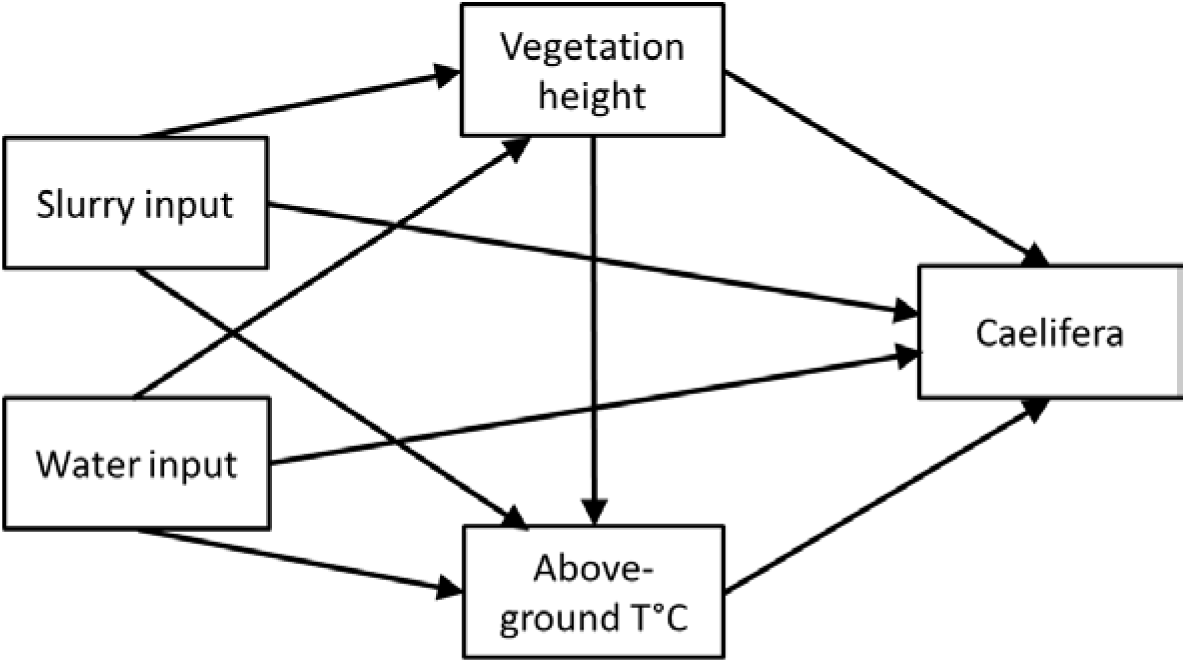

## Appendix 3.

Results of the GLMMs carried out on the effects of fertilisation and irrigation on Caelifera and Ensifera densities for both sampling sessions (i.e. before and after mowing). Table refers to figure 1 in the article. The fixed factors were the experimental treatments (C = control plots; F = fertilised; I = irrigated; I+F 1/3 = irrigation and fertilisation at low dose; I+F 2/3 = irrigation and fertilisation at medium dose; I+F 3/3 = irrigation and fertilisation at high dose). The random factors were the study sites. Parameter estimates (differences between expected mean abundances on the log scale) are given for paired regime comparisons and significant differences are highlighted in bold.

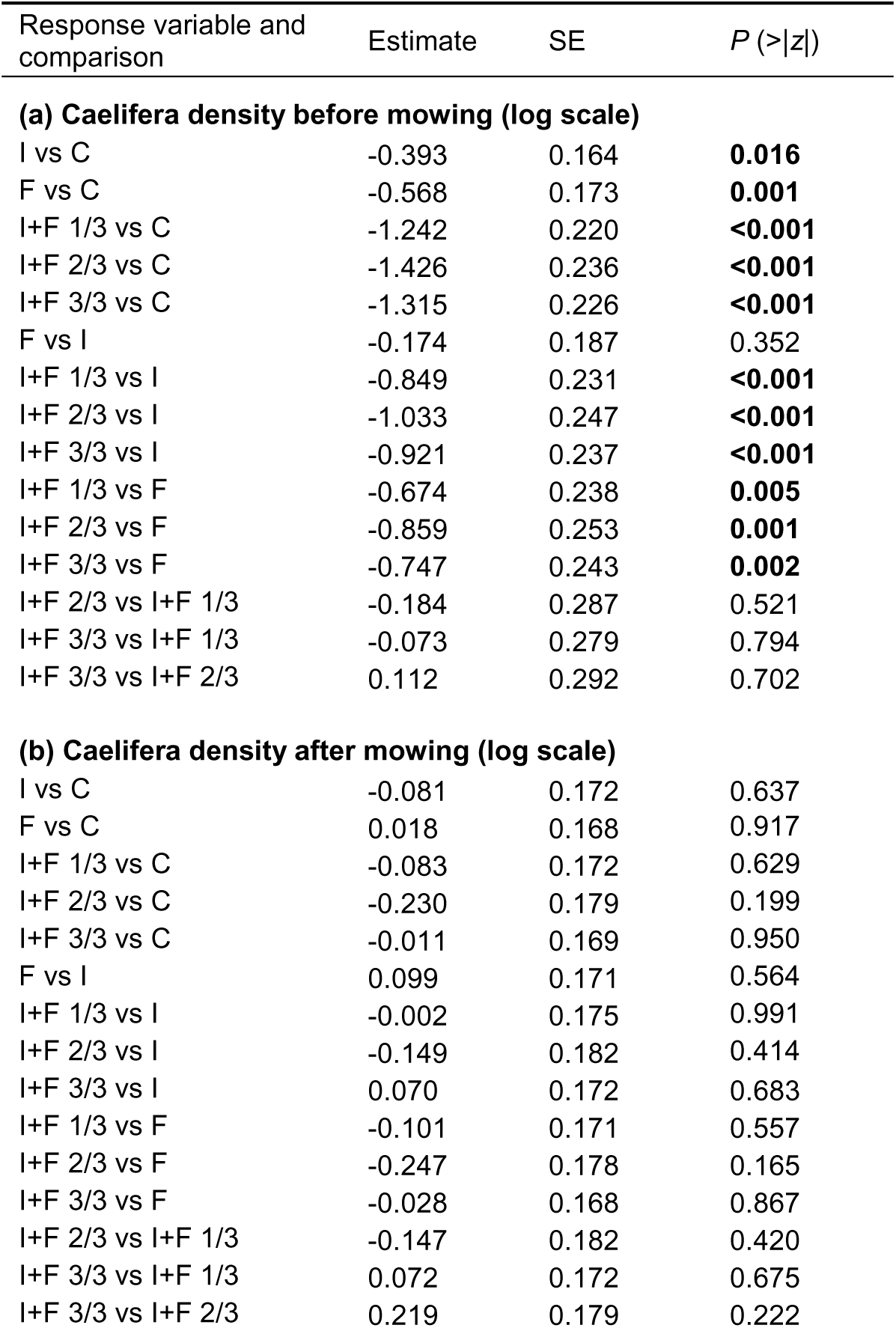
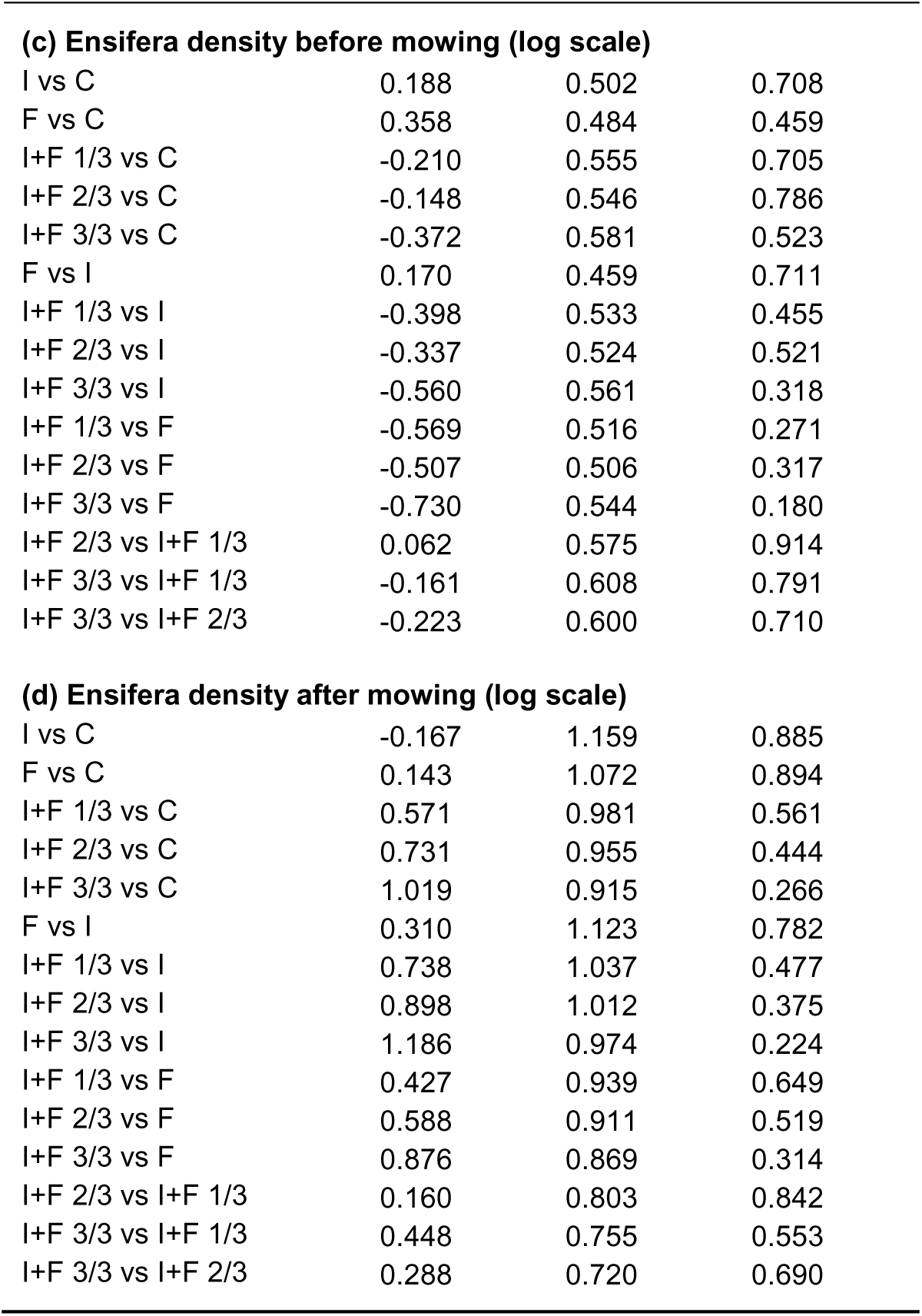

| Response variable and comparison | Estimate | SE | *P* (>\|z\|) |
| --- | --- | --- | --- |
| **(a) Caelifera density before mowing (log scale)** |  |  |  |
| I vs C | -0.393 | 0.164 | **0.016** |
| F vs C | -0.568 | 0.173 | **0.001** |
| I+F 1/3 vs C | -1.242 | 0.220 | **<0.001** |
| I+F 2/3 vs C | -1.426 | 0.236 | **<0.001** |
| I+F 3/3 vs C | -1.315 | 0.226 | **<0.001** |
| F vs I | -0.174 | 0.187 | 0.352 |
| I+F 1/3 vs I | -0.849 | 0.231 | **<0.001** |
| I+F 2/3 vs I | -1.033 | 0.247 | **<0.001** |
| I+F 3/3 vs I | -0.921 | 0.237 | **<0.001** |
| I+F 1/3 vs F | -0.674 | 0.238 | **0.005** |
| I+F 2/3 vs F | -0.859 | 0.253 | **0.001** |
| I+F 3/3 vs F | -0.747 | 0.243 | **0.002** |
| I+F 2/3 vs I+F 1/3 | -0.184 | 0.287 | 0.521 |
| I+F 3/3 vs I+F 1/3 | -0.073 | 0.279 | 0.794 |
| I+F 3/3 vs I+F 2/3 | 0.112 | 0.292 | 0.702 |
| **(b) Caelifera density after mowing (log scale)** |  |  |  |
| I vs C | -0.081 | 0.172 | 0.637 |
| F vs C | 0.018 | 0.168 | 0.917 |
| I+F 1/3 vs C | -0.083 | 0.172 | 0.629 |
| I+F 2/3 vs C | -0.230 | 0.179 | 0.199 |
| I+F 3/3 vs C | -0.011 | 0.169 | 0.950 |
| F vs I | 0.099 | 0.171 | 0.564 |
| I+F 1/3 vs I | -0.002 | 0.175 | 0.991 |
| I+F 2/3 vs I | -0.149 | 0.182 | 0.414 |
| I+F 3/3 vs I | 0.070 | 0.172 | 0.683 |
| I+F 1/3 vs F | -0.101 | 0.171 | 0.557 |
| I+F 2/3 vs F | -0.247 | 0.178 | 0.165 |
| I+F 3/3 vs F | -0.028 | 0.168 | 0.867 |
| I+F 2/3 vs I+F 1/3 | -0.147 | 0.182 | 0.420 |
| I+F 3/3 vs I+F 1/3 | 0.072 | 0.172 | 0.675 |
| I+F 3/3 vs I+F 2/3 | 0.219 | 0.179 | 0.222 |
| **(c) Ensifera density before mowing (log scale)** |  |  |  |
| I vs C | 0.188 | 0.502 | 0.708 |
| F vs C | 0.358 | 0.484 | 0.459 |
| I+F 1/3 vs C | -0.210 | 0.555 | 0.705 |
| I+F 2/3 vs C | -0.148 | 0.546 | 0.786 |
| I+F 3/3 vs C | -0.372 | 0.581 | 0.523 |
| F vs I | 0.170 | 0.459 | 0.711 |
| I+F 1/3 vs I | -0.398 | 0.533 | 0.455 |
| I+F 2/3 vs I | -0.337 | 0.524 | 0.521 |
| I+F 3/3 vs I | -0.560 | 0.561 | 0.318 |
| I+F 1/3 vs F | -0.569 | 0.516 | 0.271 |
| I+F 2/3 vs F | -0.507 | 0.506 | 0.317 |
| I+F 3/3 vs F | -0.730 | 0.544 | 0.180 |
| I+F 2/3 vs I+F 1/3 | 0.062 | 0.575 | 0.914 |
| I+F 3/3 vs I+F 1/3 | -0.161 | 0.608 | 0.791 |
| I+F 3/3 vs I+F 2/3 | -0.223 | 0.600 | 0.710 |
| **(d) Ensifera density after mowing (log scale)** |  |  |  |
| I vs C | -0.167 | 1.159 | 0.885 |
| F vs C | 0.143 | 1.072 | 0.894 |
| I+F 1/3 vs C | 0.571 | 0.981 | 0.561 |
| I+F 2/3 vs C | 0.731 | 0.955 | 0.444 |
| I+F 3/3 vs C | 1.019 | 0.915 | 0.266 |
| F vs I | 0.310 | 1.123 | 0.782 |
| I+F 1/3 vs I | 0.738 | 1.037 | 0.477 |
| I+F 2/3 vs I | 0.898 | 1.012 | 0.375 |
| I+F 3/3 vs I | 1.186 | 0.974 | 0.224 |
| I+F 1/3 vs F | 0.427 | 0.939 | 0.649 |
| I+F 2/3 vs F | 0.588 | 0.911 | 0.519 |
| I+F 3/3 vs F | 0.876 | 0.869 | 0.314 |
| I+F 2/3 vs I+F 1/3 | 0.160 | 0.803 | 0.842 |
| I+F 3/3 vs I+F 1/3 | 0.448 | 0.755 | 0.553 |
| I+F 3/3 vs I+F 2/3 | 0.288 | 0.720 | 0.690 |

## Appendix 4.

Results of the LMMs carried out on the effects of fertilisation and irrigation on Caelifera and Ensifera species richness. Table refers to figure 2 in the article. Both sampling sessions were analysed together. The fixed factors were the experimental treatments (C = control plots; F = fertilised; I = irrigated; I+F 1/3 = irrigation and fertilisation at low dose; I+F 2/3 = irrigation and fertilisation at medium dose; I+F 3/3 = irrigation and fertilisation at high dose). The random factors were the study sites. Parameter estimates (differences between expected mean species richness) are given for paired regime comparisons and significant differences are highlighted in bold.

| Response variable and comparison | Estimate | SE | *P* (>\|t\|) |
| --- | --- | --- | --- |
| **(a) Caelifera species richness** |  |  |  |
| I vs C | -0.333 | 0.358 | 0.355 |
| F vs C | -0.417 | 0.358 | 0.249 |
| I+F 1/3 vs C | -1.417 | 0.358 | **<0.001** |
| I+F 2/3 vs C | -1.083 | 0.358 | **0.004** |
| I+F 3/3 vs C | -2.250 | 0.358 | **<0.001** |
| F vs I | -0.083 | 0.357 | 0.817 |
| I+F 1/3 vs I | -1.083 | 0.357 | **0.004** |
| I+F 2/3 vs I | -0.750 | 0.357 | **0.041** |
| I+F 3/3 vs I | -1.917 | 0.357 | **<0.001** |
| I+F 1/3 vs F | -1.000 | 0.357 | **0.007** |
| I+F 2/3 vs F | -0.667 | 0.357 | 0.068 |
| I+F 3/3 vs F | -1.833 | 0.357 | **<0.001** |
| I+F 2/3 vs I+F 1/3 | 0.333 | 0.358 | 0.355 |
| I+F 3/3 vs I+F 1/3 | -0.833 | 0.358 | **0.023** |
| I+F 3/3 vs I+F 2/3 | -1.167 | 0.358 | **0.002** |
| **(b) Ensifera species richness** |  |  |  |
| I vs C | -0.083 | 0.332 | 0.803 |
| F vs C | 0.167 | 0.332 | 0.618 |
| I+F 1/3 vs C | 0.417 | 0.332 | 0.215 |
| I+F 2/3 vs C | 0.250 | 0.332 | 0.455 |
| I+F 3/3 vs C | 0.417 | 0.332 | 0.215 |
| F vs I | 0.250 | 0.332 | 0.455 |
| I+F 1/3 vs I | 0.500 | 0.332 | 0.138 |
| I+F 2/3 vs I | 0.333 | 0.332 | 0.320 |
| I+F 3/3 vs I | 0.500 | 0.332 | 0.138 |
| I+F 1/3 vs F | 0.250 | 0.332 | 0.455 |
| I+F 2/3 vs F | 0.083 | 0.332 | 0.803 |
| I+F 3/3 vs F | 0.250 | 0.332 | 0.455 |
| I+F 2/3 vs I+F 1/3 | -0.167 | 0.332 | 0.618 |
| I+F 3/3 vs I+F 1/3 | 0.000 | 0.332 | 1.000 |
| I+F 3/3 vs I+F 2/3 | 0.167 | 0.332 | 0.618 |

## Appendix 5.

Results of the LMMs carried out on the effects of fertilisation and irrigation on average vegetation height for both sampling sessions (i.e. before and after mowing). Table refers to figure 3 in the article. The fixed factors were the experimental treatments (C = control plots; F = fertilised; I = irrigated; I+F 1/3 = irrigation and fertilisation at low dose; I+F 2/3 = irrigation and fertilisation medium at dose; I+F 3/3 = irrigation and fertilisation at high dose). The random factors were the study sites. Parameter estimates (differences between expected mean) are given for paired regime comparisons and significant differences are highlighted in bold.

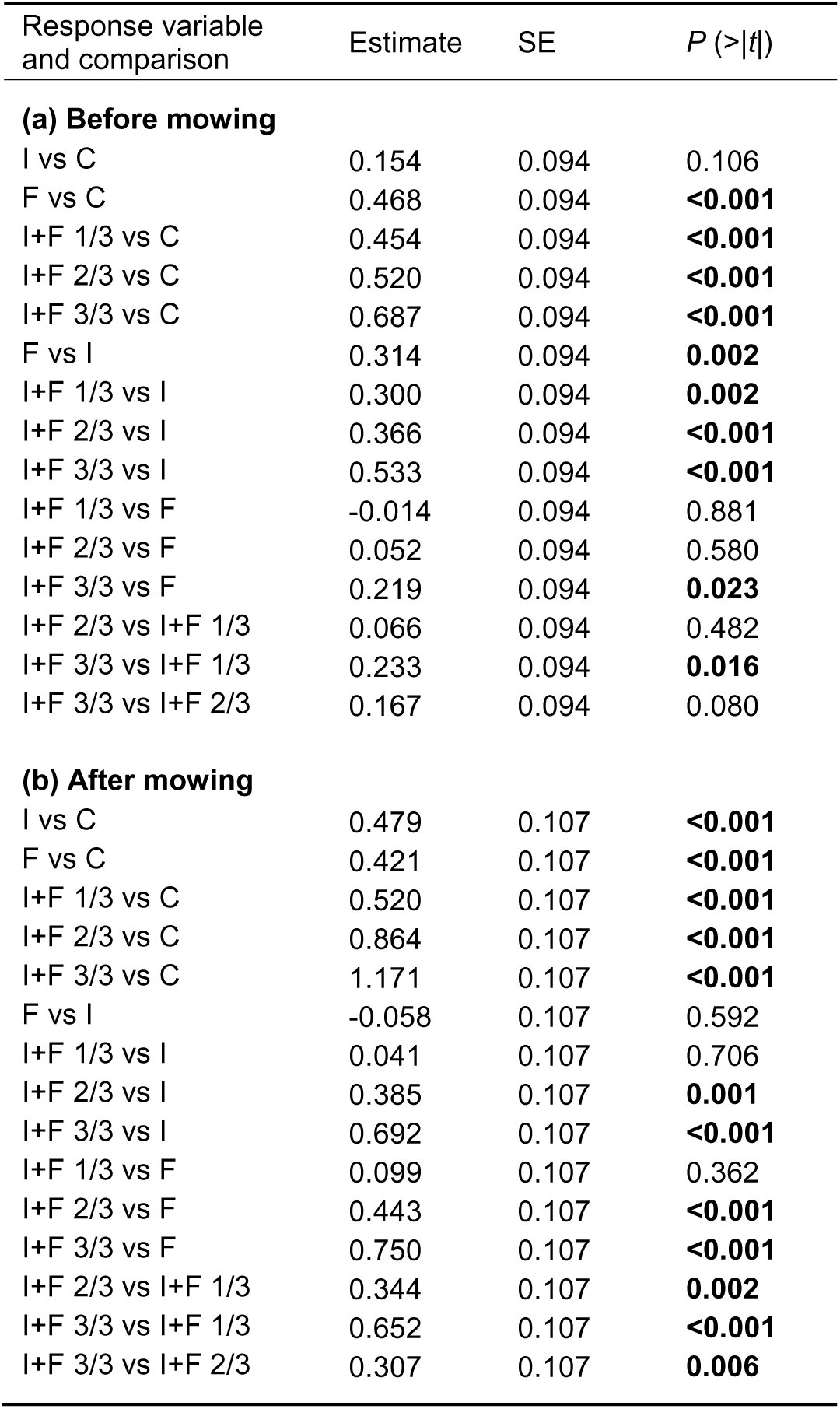

| Response variable and comparison | Estimate | SE | *P* (>\|*t*\|) |
| --- | --- | --- | --- |
| **(a) Before mowing** |  |  |  |
| I vs C | 0.154 | 0.094 | 0.106 |
| F vs C | 0.468 | 0.094 | **<0.001** |
| I+F 1/3 vs C | 0.454 | 0.094 | **<0.001** |
| I+F 2/3 vs C | 0.520 | 0.094 | **<0.001** |
| I+F 3/3 vs C | 0.687 | 0.094 | **<0.001** |
| F vs I | 0.314 | 0.094 | **0.002** |
| I+F 1/3 vs I | 0.300 | 0.094 | **0.002** |
| I+F 2/3 vs I | 0.366 | 0.094 | **<0.001** |
| I+F 3/3 vs I | 0.533 | 0.094 | **<0.001** |
| I+F 1/3 vs F | -0.014 | 0.094 | 0.881 |
| I+F 2/3 vs F | 0.052 | 0.094 | 0.580 |
| I+F 3/3 vs F | 0.219 | 0.094 | **0.023** |
| I+F 2/3 vs I+F 1/3 | 0.066 | 0.094 | 0.482 |
| I+F 3/3 vs I+F 1/3 | 0.233 | 0.094 | **0.016** |
| I+F 3/3 vs I+F 2/3 | 0.167 | 0.094 | 0.080 |
| **(b) After mowing** |  |  |  |
| I vs C | 0.479 | 0.107 | **<0.001** |
| F vs C | 0.421 | 0.107 | **<0.001** |
| I+F 1/3 vs C | 0.520 | 0.107 | **<0.001** |
| I+F 2/3 vs C | 0.864 | 0.107 | **<0.001** |
| I+F 3/3 vs C | 1.171 | 0.107 | **<0.001** |
| F vs I | -0.058 | 0.107 | 0.592 |
| I+F 1/3 vs I | 0.041 | 0.107 | 0.706 |
| I+F 2/3 vs I | 0.385 | 0.107 | **0.001** |
| I+F 3/3 vs I | 0.692 | 0.107 | **<0.001** |
| I+F 1/3 vs F | 0.099 | 0.107 | 0.362 |
| I+F 2/3 vs F | 0.443 | 0.107 | **<0.001** |
| I+F 3/3 vs F | 0.750 | 0.107 | **<0.001** |
| I+F 2/3 vs I+F 1/3 | 0.344 | 0.107 | **0.002** |
| I+F 3/3 vs I+F 1/3 | 0.652 | 0.107 | **<0.001** |
| I+F 3/3 vs I+F 2/3 | 0.307 | 0.107 | **0.006** |

## Appendix 6.

Results of the LMMs carried out on the effects of fertilisation and irrigation on aboveground diurnal temperature for both sampling sessions (i.e. before and after mowing). Table refers to figure 3 in the article. The fixed factors were the experimental treatments (C = control plots; F = fertilised; I = irrigated; I+F 1/3 = irrigation and fertilisation at low dose; I+F 2/3 = irrigation and fertilisation medium at dose; I+F 3/3 = irrigation and fertilisation at high dose). The random factors were the study sites. Parameter estimates (differences between expected mean) are given for paired regime comparisons and significant differences are highlighted in bold.

| Response variable and comparison | Estimate | SE | *P* (>\|*t*\|) |
| --- | --- | --- | --- |
| **(a) Before mowing** |  |  |  |
| I vs C | -1.300 | 0.850 | 0.134 |
| F vs C | -2.111 | 0.920 | **0.027** |
| I+F 1/3 vs C | -2.305 | 1.098 | **0.042** |
| I+F 2/3 vs C | -2.099 | 0.876 | **0.021** |
| I+F 3/3 vs C | -4.317 | 0.866 | **<0.001** |
| F vs I | -0.810 | 0.914 | 0.381 |
| I+F 1/3 vs I | -1.005 | 1.081 | 0.358 |
| I+F 2/3 vs I | -0.799 | 0.849 | 0.353 |
| I+F 3/3 vs I | -3.017 | 0.849 | **0.001** |
| I+F 1/3 vs F | -0.195 | 1.133 | 0.865 |
| I+F 2/3 vs F | 0.012 | 0.929 | 0.990 |
| I+F 3/3 vs F | -2.207 | 0.930 | **0.023** |
| I+F 2/3 vs I+F 1/3 | 0.206 | 1.094 | 0.851 |
| I+F 3/3 vs I+F 1/3 | -2.012 | 1.099 | 0.075 |
| I+F 3/3 vs I+F 2/3 | -2.218 | 0.875 | **0.015** |
| **(b) After mowing** |  |  |  |
| I vs C | -1.612 | 0.949 | 0.097 |
| F vs C | 0.181 | 0.949 | 0.850 |
| I+F 1/3 vs C | -0.503 | 0.949 | 0.599 |
| I+F 2/3 vs C | -1.636 | 0.948 | 0.092 |
| I+F 3/3 vs C | -1.228 | 1.028 | 0.240 |
| F vs I | 1.793 | 0.979 | 0.075 |
| I+F 1/3 vs I | 1.109 | 0.968 | 0.259 |
| I+F 2/3 vs I | -0.024 | 0.978 | 0.981 |
| I+F 3/3 vs I | 0.385 | 1.060 | 0.719 |
| I+F 1/3 vs F | -0.684 | 0.979 | 0.489 |
| I+F 2/3 vs F | -1.817 | 0.978 | 0.071 |
| I+F 3/3 vs F | -1.408 | 1.046 | 0.186 |
| I+F 2/3 vs I+F 1/3 | -1.133 | 0.978 | 0.254 |
| I+F 3/3 vs I+F 1/3 | -0.724 | 1.060 | 0.499 |
| I+F 3/3 vs I+F 2/3 | 0.409 | 1.058 | 0.702 |

