## Appendix 1 for "Grassland irrigation and fertilisation alter vegetation height and vegetation within temperature and negatively affect orthopteran populations"

**Appendix 1.** Location name of all study sites (n = 12), geographic coordinates (WGS84), altitude, sampling date, and number of orthopterans of each species caught during the two sampling sessions (i.e. before and after mowing). In addition, meadows were classified in three groups according to the maximum hay productivity potential of the site (see Material and methods section for more details). Data are missing for the first session in Cordona due to technical problems.

| | Site | Coordinates | | Group | Altitude [m] | Sampling date | Caelifera larvae | Ensifera larvae | *Pseudochorthippus parallelus* | *Chortippus biguttulus* | *Chortippus dorsatus* | *Chortippus brunneus* | *Stauroderus scalaris* | *Stenobothrus lineatus* | *Euthystira brachyptera* | *Omocestus rufipes* | *Omocestus virudulus* | *Omocestus haemorrhoidalis* | *Acryptera fusca* | *Chrysochraon dispar* | *Mecosesthus parapleurus* | *Tettigonia viridissima* | *Roeseliana roeselii* | *Metrioptera saussuriana* | *Decticus verrucivorus* | *Plactycleis albopunctata* | *Pholidoptera griseoaptera* | *Conocephalus fuscus* | *Leptophyes punctatissima* |
|---|---|---|---|---|---|---|---|---|---|---|---|---|---|---|---|---|---|---|---|---|---|---|---|---|---|---|---|---|---|
| | | Latitude | Longitude | | | | | | | | | | | | | | | | | | | | | | | | | | |
| Before mowing | Euseigne | 46°10′9″N | 7°25′27″E | 1 | 1028 | 02.07.2013 | 36 | 24 | 0 | 0 | 0 | 0 | 0 | 0 | 0 | 0 | 0 | 0 | 0 | 0 | 0 | 0 | 0 | 0 | 0 | 0 | 0 | 0 | 0 |
| | Icogne 1 | 46°17′56″N | 7°26′31″E | 1 | 880 | 13.06.2013 | 56 | 41 | 0 | 0 | 0 | 0 | 0 | 0 | 0 | 0 | 0 | 0 | 0 | 0 | 0 | 0 | 0 | 0 | 4 | 0 | 0 | 0 | 0 |
| | Orsieres | 46°1′44″N | 7°9′8″E | 1 | 1022 | 19.06.2013 | 90 | 15 | 0 | 0 | 0 | 0 | 0 | 0 | 0 | 0 | 0 | 0 | 0 | 0 | 0 | 0 | 0 | 0 | 2 | 0 | 0 | 0 | 0 |
| | Sembrancher | 46°4′24″N | 7°8′36″E | 1 | 798 | 12.06.2013 | 27 | 15 | 0 | 0 | 0 | 0 | 0 | 0 | 0 | 0 | 0 | 0 | 0 | 0 | 0 | 0 | 0 | 0 | 0 | 0 | 0 | 0 | 0 |
| | Arbaz | 46°16′42″N | 7°22′47″E | 2 | 1270 | 26.06.2013 | 27 | 22 | 0 | 0 | 0 | 0 | 0 | 0 | 7 | 0 | 0 | 0 | 0 | 0 | 0 | 0 | 0 | 0 | 3 | 0 | 0 | 0 | 0 |
| | Cordona | 46°19′45″N | 7°33′8″E | 2 | 1153 | 12.06.2013 | NA | NA | NA | NA | NA | NA | NA | NA | NA | NA | NA | NA | NA | NA | NA | NA | NA | NA | NA | NA | NA | NA | NA |
| | Icogne 2 | 46°16′42″N | 7°26′10″E | 2 | 1200 | 26.06.2013 | 48 | 40 | 0 | 0 | 0 | 0 | 0 | 0 | 0 | 0 | 0 | 0 | 0 | 0 | 0 | 0 | 0 | 0 | 8 | 0 | 0 | 0 | 0 |
| | La Garde | 46°3′45″N | 7°8′35″E | 2 | 980 | 18.06.2013 | 267 | 73 | 0 | 1 | 0 | 0 | 0 | 0 | 0 | 0 | 0 | 0 | 0 | 0 | 0 | 0 | 0 | 0 | 14 | 2 | 0 | 0 | 0 |
| | Vens | 46°5′7″N | 7°7′24″E | 2 | 1373 | 12.06.2013 | 119 | 29 | 0 | 0 | 0 | 0 | 0 | 0 | 0 | 0 | 0 | 0 | 0 | 0 | 0 | 0 | 0 | 0 | 0 | 0 | 0 | 0 | 0 |
| | Eison | 46°9′18″N | 7°28′10″E | 3 | 1768 | 12.07.2013 | 708 | 29 | 0 | 0 | 0 | 0 | 0 | 0 | 15 | 0 | 0 | 0 | 0 | 0 | 0 | 0 | 0 | 0 | 4 | 0 | 0 | 0 | 0 |
| | Grimentz | 46°11′22″N | 7°34′35″E | 3 | 1738 | 08.07.2013 | 360 | 6 | 0 | 1 | 0 | 0 | 0 | 1 | 0 | 0 | 1 | 0 | 0 | 0 | 0 | 0 | 0 | 0 | 0 | 0 | 0 | 0 | 0 |
| | St-Martin | 46°11′8″N | 7°26′43″E | 3 | 1589 | 02.07.2013 | 488 | 17 | 0 | 0 | 0 | 0 | 0 | 0 | 0 | 0 | 0 | 0 | 0 | 0 | 0 | 0 | 0 | 0 | 0 | 0 | 0 | 0 | 0 |
| After mowing | Euseigne | 46°10′9″N | 7°25′27″E | 1 | 1028 | 22.08.2013 | 30 | 0 | 39 | 3 | 42 | 7 | 0 | 5 | 1 | 0 | 0 | 0 | 4 | 0 | 0 | 1 | 2 | 3 | 0 | 0 | 1 | 0 | 0 |
| | Icogne 1 | 46°17′56″N | 7°26′31″E | 1 | 880 | 14.08.2013 | 39 | 0 | 7 | 0 | 0 | 0 | 8 | 4 | 2 | 2 | 0 | 0 | 0 | 1 | 0 | 1 | 3 | 0 | 0 | 1 | 1 | 0 | 0 |
| | Orsieres | 46°1′44″N | 7°9′8″E | 1 | 1022 | 21.08.2013 | 63 | 3 | 29 | 22 | 1 | 0 | 3 | 1 | 11 | 0 | 0 | 1 | 0 | 0 | 4 | 1 | 4 | 0 | 0 | 0 | 0 | 2 | 0 |
| | Sembrancher | 46°4′24″N | 7°8′36″E | 1 | 798 | 13.08.2013 | 17 | 0 | 33 | 0 | 5 | 0 | 3 | 3 | 7 | 0 | 0 | 0 | 0 | 0 | 0 | 1 | 2 | 0 | 0 | 0 | 0 | 0 | 0 |
| | Arbaz | 46°16′42″N | 7°22′47″E | 2 | 1270 | 23.08.2013 | 190 | 0 | 2 | 11 | 0 | 0 | 1 | 1 | 5 | 0 | 0 | 0 | 0 | 0 | 0 | 0 | 0 | 0 | 2 | 3 | 0 | 0 | 0 |
| | Cordona | 46°19′45″N | 7°33′8″E | 2 | 1153 | 23.08.3013 | 280 | 1 | 305 | 15 | 10 | 4 | 7 | 6 | 2 | 2 | 0 | 1 | 0 | 0 | 1 | 2 | 15 | 0 | 3 | 0 | 0 | 0 | 8 |
| | Icogne 2 | 46°16′42″N | 7°26′10″E | 2 | 1200 | 14.08.2013 | 419 | 0 | 36 | 0 | 0 | 0 | 9 | 1 | 0 | 0 | 0 | 0 | 0 | 0 | 0 | 0 | 4 | 0 | 0 | 0 | 0 | 0 | 0 |
| | La Garde | 46°3′45″N | 7°8′35″E | 2 | 980 | 13.08.2013 | 20 | 0 | 48 | 2 | 2 | 0 | 52 | 8 | 3 | 0 | 0 | 5 | 0 | 0 | 0 | 1 | 4 | 1 | 6 | 10 | 9 | 0 | 0 |
| | Vens | 46°5′7″N | 7°7′24″E | 2 | 1373 | 21.08.2013 | 36 | 0 | 0 | 8 | 0 | 4 | 13 | 2 | 0 | 5 | 0 | 0 | 4 | 0 | 0 | 0 | 6 | 0 | 0 | 0 | 1 | 0 | 0 |
| | Eison | 46°9′18″N | 7°28′10″E | 3 | 1768 | 31.08.2013 | 477 | 0 | 191 | 12 | 24 | 0 | 116 | 117 | 18 | 0 | 0 | 26 | 2 | 0 | 0 | 1 | 3 | 3 | 0 | 0 | 0 | 0 | 0 |
| | Grimentz | 46°11′22″N | 7°34′35″E | 3 | 1738 | 26.08.2013 | 35 | 0 | 13 | 2 | 16 | 3 | 46 | 1 | 0 | 8 | 0 | 0 | 0 | 0 | 0 | 0 | 0 | 0 | 0 | 0 | 0 | 0 | 0 |
| | St-Martin | 46°11′8″N | 7°26′43″E | 3 | 1589 | 31.08.2013 | 27 | 0 | 6 | 3 | 0 | 0 | 59 | 0 | 6 | 1 | 0 | 0 | 6 | 0 | 0 | 0 | 7 | 4 | 3 | 0 | 0 | 0 | 0 |
